# Characterizing dry mass and volume changes in human multiple myeloma cells upon treatment with proteotoxic and genotoxic drugs

**DOI:** 10.1101/2022.03.18.484926

**Authors:** Xili Liu, Maria Moscvin, Seungeun Oh, Tianzeng Chen, Wonshik Choi, Benjamin Evans, Sean M. Rowell, Omar Nadeem, Clifton C. Mo, Adam S. Sperling, Kenneth C. Anderson, Zahid Yaqoob, Giada Bianchi, Yongjin Sung

## Abstract

Multiple myeloma (MM) is a cancer of terminally differentiated plasma cells. MM remains incurable, but overall survival of patients has progressively increased over the past two decades largely due to novel agents such as proteasome inhibitors (PI) and the immunomodulatory agents. While these therapies are highly effective, MM patients can be *de novo* resistant and acquired resistance with prolonged treatment is inevitable. There is growing interest in early, accurate identification of responsive versus non-responsive patients, however limited sample availability and need for rapid assays are limiting factors. Here, we test dry mass and volume as label-free biomarkers to monitor early response of MM cells to treatment with bortezomib, doxorubicin, and ultraviolet light. For the dry mass measurement, we use two types of phase-sensitive optical microscopy techniques: digital holographic tomography and computationally-enhanced quantitative phase microscopy. We show that human MM cell lines (RPMI8226, MM.1S, KMS20, and AMO1) increase dry mass upon bortezomib treatment. This dry mass increase after bortezomib treatment occurs as early as 1 hour for sensitive cells and 4 hours for all tested cells. We further confirm this observation using primary multiple myeloma cells derived from patients and show that a correlation exists between increase in dry mass and sensitivity to bortezomib, supporting the use of dry mass as a biomarker. The volume measurement using Coulter counter shows a more complex behavior; RPMI8226 cells increase the volume at an early stage of apoptosis, but MM.1S cells show the volume decrease typically observed with apoptotic cells. Altogether, this cell study presents complex kinetics of dry mass and volume at an early stage of apoptosis, which may serve as a basis for the detection and treatment of MM cells.

## Introduction

Multiple myeloma (MM) is a cancer of terminally-differentiated, antibody-producing plasma cells^1^. It represents the second most common hematologic malignancy in the USA, accounting for about 18% of all hematologic tumors^2^. The introduction of novel therapies has substantially prolonged the median overall survival of MM patients from 2-3 years to 7-8 years, but development of resistance is essentially inevitable for most patients overtime, leading to patient demise^3^. In the phase III clinical trial of single agent bortezomib versus dexamethasone in relapsed/refractory MM, 43% of patients had a response to bortezomib at 22 months follow up with 9% patients achieving a complete response^4^. While these figures were impressive for a disease that had not seen any drug development since the introduction of melphalan and prednisone in the 1960s, they also outlined that over 50% of MM patients are *de novo* resistant to single agent bortezomib. It is important to point out that during the induction phase of MM treatment, multi-drug, combinatorial approaches are standard of care and single agent therapy is reserved for the maintenance phase of treatment, after adequate debulking and remission state have been achieved. The mechanisms behind bortezomib resistance remain elusive and no predictive factor of response is available for clinical use^5^. We previously showed that an imbalance between proteasome capacity and load (i.e., proteins in cue for proteasomal degradation) is predictive of bortezomib sensitivity in cell lines and primary cells from patients^6,7^. Here, using four human MM cell lines (RPMI8226, MM.1S, KMS20, and AMO1) with distinct genetic characteristics and baseline sensitivity to proteasome inhibitors, we test cell dry mass and volume as potential biomarkers to predict the sensitivity of MM cells to bortezomib, doxorubicin, and UV irradiation. Cell dry mass is a direct measure of the balance between mass production and disposal; thus, we hypothesize that the accumulation of unneeded or damaged proteins will increase the cell dry mass, which may be detectable with new highly sensitive methods. Since cells change their volume to maintain intracellular homeostasis, we hypothesized that the mass imbalance may result in an increase in volume. To measure the dry mass, we used a well-established linear relationship between the refractive index and the density of cellular materials^8,9^. By using digital holographic tomography (DHT), we can measure the 3D refractive index distribution within a heterogeneous biological specimen; thereby, it can accurately measure the dry mass of cells irrespective of the sample thickness^10^. For the dry mass measurement of a large number of cells, we also used high-throughput, computationally-enhanced quantitative phase microscopy (ceQPM), which records the projected refractive index map^11^. To measure the volume, we used the Coulter counter, which measures the volume of a cell using the change in electrical impedance when the cell traverses a small orifice^12^. These single-cell, physical measurements can also be applied to primary cells without any special sample preparation except for standard gradient centrifugation and magnetic bead-mediated positive selection.

## Materials and Methods

### Cell culture

All human MM cell lines (RPMI8226, MM.1S, KMS20, and AMO1) were purchased from ATCC. Bortezomib resistant AMO1 cell line (AMO1-VR) was a kind gift of Dr. Christopher Driessen, Kantonsspital St. Gallen, Switzerland. All MM cell lines were maintained in Roswell Park Memorial Institute (RPMI)-1640 medium containing 2.5 mg/mL plasmocin, 1x Gibco Antibiotic-Antimycotic (100 U/mL of penicillin, 100 μg/mL of streptomycin, and 25 ng/mL of Gibco Amphotericin B), supplemented with 10% (v/v) fetal bovine serum (FBS) and 2 μg L-glutamine. HeLa human cervical cancer cells were purchased from ATCC, cultured in Dulbecco’s modified Eagle’s medium (Invitrogen, 21063-029) supplemented with 10% fetal bovine serum and 1% 100X penicillin-streptomycin solution. All cell lines were authenticated via All cell lines were authenticated via short tandem repeat (STR) profiling performed by the Dana Farber Cancer Institute Molecular Diagnostic Laboratory. Mycoplasma testing was routinely performed prior to experiments.

### Primary MM cells

Bone marrow aspirate samples were obtained from patients newly diagnosed or with relapsed/refractory MM treated at Dana Farber Cancer Institute. Informed consent was obtained from all participants and/or their legal guardians according to a research protocol approval by the Institutional Review Board of the Dana-Farber Cancer Institute. All research activities were performed in accordance with relevant instutional guidelines and regulations and in accordance with the [Declaration of Helsinki](https://www.wma.net/policies-post/wma-declaration-of-helsinki-ethical-principles-for-medical-research-involving-human-subjects/). after obtaining informed consent and. Bone marrow mononuclear cells (BMMC) were separated by Ficoll-Paque PLUS (GE Healthcare), and MM cells were enriched by CD138-positive selection with magnetic activated cell separation microbeads (Miltenyi Biotec). MM cells were cultured in RPMI containing 100 U/ml penicillin and 100 ug/ml streptomycin, supplemented with 20% (v/v) FBS.

### Drugs and antibodies

Bortezomib and doxorubicin were purchased from Selleckchem as dry powder and resuspended as directed in DMSO. Just prior to use in experiments, bortezomib and doxorubicin were resuspended to a working concentration in RPMI media. Antibodies used in this study as follows: PARP (Cell Signaling, 9532); GAPDH (Cell Signaling, 5174); K48-linked ubiquitin (Apu2 clone, Sigma-Aldrich, 05-1307). As secondary antibody we used anti rabbit IgG, HRP conjugated from Cell Signaling.

### Cell dry mass measurement using digital holographic tomography (DHT) and computationally enhanced quantitative phase microscopy (ceQPM)

Cell dry mass can be measured from refractive index (*n*), which is proportional to the concentration (C) of cellular dry mass as *n* = *n*_0_ + *αC*, where *n*_0_ is the refractive index of the culture medium. The specific refractive index increment, α, is constant at 0.190 mL/g, regardless of the chemical identity of the biomaterial^8,9^. Thus, the cell dry mass (*m*) can be calculated using Eq. (1).

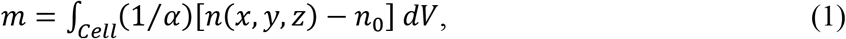

where *n*(*x, y, z*) is the 3D map of refractive index for a cell. In the present work, we use DHT^13–16^ to measure the 3D refractive index map of a cell (Fig. 1A), from which the dry mass is calculated using Eq. (1). Figure 1B shows example cross-sections of the 3D refractive index maps of an RPMI8226 cell before and 4 hours of treatment with 20 nM bortezomib. For the DHT measurement, cells were harvested at 70 - 80% confluency using Trypsin-EDTA (Invitrogen, 25200), seeded on a glass cover slip precoated with poly-L-lysine (P8920; Sigma–Aldrich). After incubation for 4 hours in a 5% CO_2_ cell culture incubator at 37 °C, the cells were treated with the indicated dose of bortezomib, or doxorubicin, or irradiated with ultraviolet (UV) light for the indicated time. At the end of treatment, cells were gently washed twice with fresh warm medium and imaged using DHT. <u>For each test case, we imaged about 70 fields-of-view (FOV) within 40 minutes. Each FOV typically contains 1-2 cells</u>.

**Figure 1.**
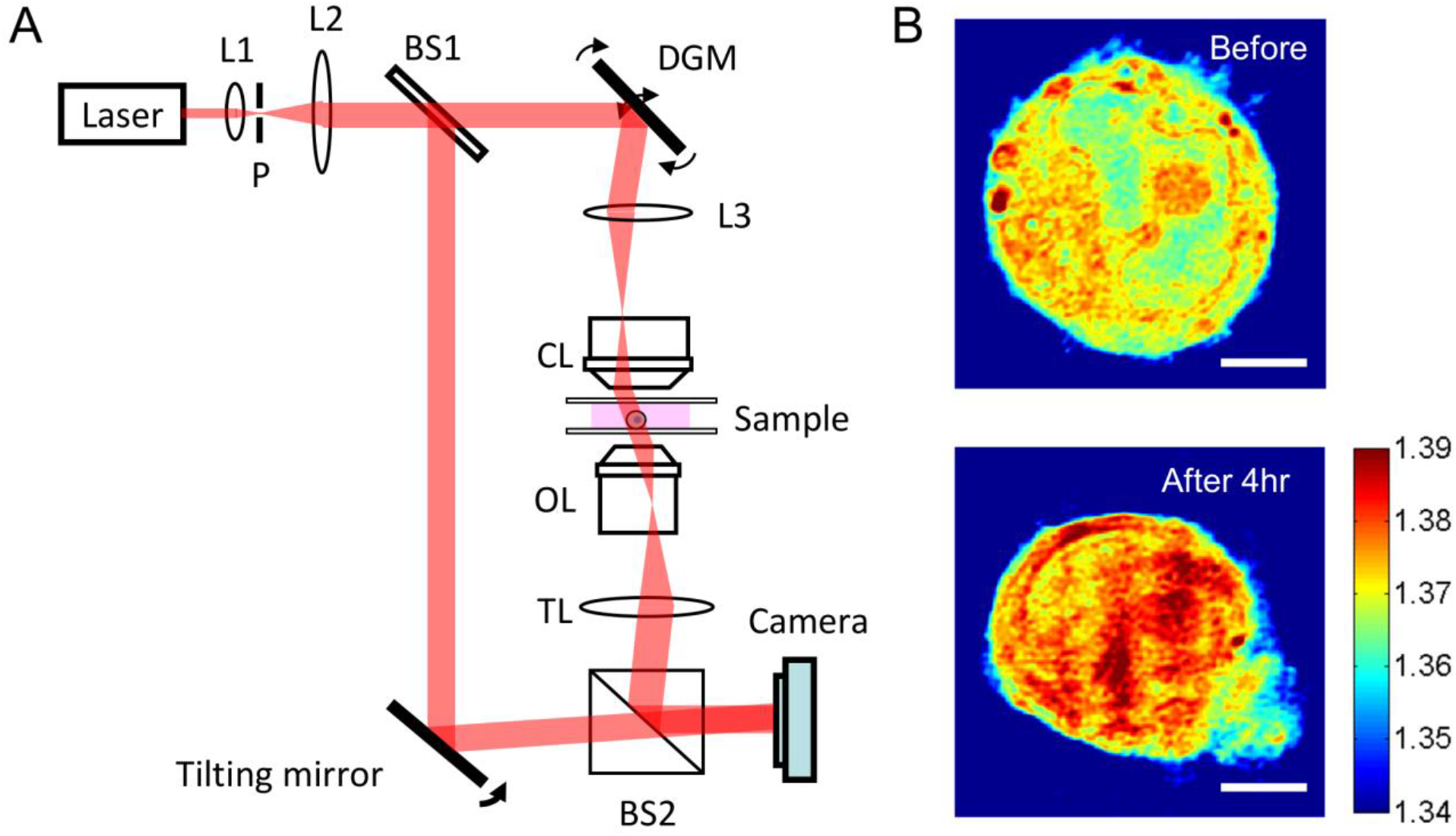
Imaging myeloma cells with digital holographic tomography. **(A)** Digital holographic tomography for 3D refractive index imaging. L: lens; M: mirror; BS: beam splitter; CL: condenser lens; OL: objective lens; TL: tube lens; DGM: dual-axis galvanometer mirror; P: pinhole. **(B)** Example refractive-index maps of RPMI cells before and after 4 hours of treatment with 20 nM bortezomib. Horizontal cross-sections of the 3D refractive index maps are shown. Scale bar: 5 μm.

For the dry mass measurement of a large number of cells, we used computationally-enhanced quantitative phase microscopy (ceQPM)^11^ with a SID4BIO (Phasics, France) camera to detect the interferogram. The principle of ceQPM is similar to DHT except for that it detects the 2D projection map of the refractive index. For ceQPM, the cultured cell lines or patient samples were treated by 16 nM bortezomib for indicated time and seeded on 6-well glass-bottom plates (Mattek, P06G-1.5-10-F) pretreated with poly-L-lysine (P8920; Sigma–Aldrich) at a density of 250 cells/mm^2^. An 18*18 glass coverslip was placed in each well to form a small chamber of filled medium which supported the cells for a brief period. The samples were imaged immediately on a Ti microscope with a Plan Apo 20× N.A. 0.75 PFS dry objective and motorized stage (Nikon, Japan). The scan of a 6-well glass-bottom plate takes less than 15 minutes. The interferograms were processed by the ceQPM algorithms that were previously developed^11^.

### Volume measurement using Coulter counter

For the volume measurement, cells were grown in 25-cm^2^ culture flasks. At 70 - 80 % confluency for adherent cells or at a concentration of circa 250,000/mL, cells were treated with bortezomib, treated with doxorubicin, or irradiated with UV light for the same dose and time as those used for the dry mass measurement. Adherent cells were harvested using 0.25% Trypsin-EDTA (Invitrogen, 25200-114) while suspension cells were harvested by pipetting. Cells were centrifuged to remove debris, and then analyzed using the Z2 Coulter Counter (Beckman Coulter, Inc.). <u>We prepared two independent samples, and for each sample, we repeated the measurement three times. The data were merged for the analysis</u>.

### Cell cycle analysis

Cells were harvested at indicated times after treatment and washed once in PBS before adding cold 80% ethanol, drop-wise to the cell pellet while vortexing. After one-hour incubation at 4 degrees, cells were washed, incubated in RNase at 37 degrees for 30 minutes before adding propidium iodide at a final concentration of 10 microg/mL in PBS. Cells were incubated overnight at 4 degrees. Cell cycle analysis was performed the following day via flow cytometry and data analyzed with FlowJo.

### Drug treatment and apoptosis assessment

MM cells were treated for the indicated dose and time with bortezomib and doxorubicin or with 60 second pulse of UV light. Cells were then either collected after the indicated treatment time and analyzed for apoptosis via flow cytometry after annexin V/propidium iodide (PI) staining according to protocol (BD Biosciences) or washed in PBS once and seeded in fresh media without drugs to complete 24 hour of incubation (wash out) at the end of which cells were harvested and analyzed for apoptosis via flow cytometry after annexin V/propidium iodide staining. Histogram bars show % of viable cells (annexin V negative, PI negative).

### Western blot

Cells were harvested as indicated before, washed once in PBS and lysed in RIPA buffer (Boston Bioproducts) containing a mixture of protease inhibitor (cOmplete, Mini, Roche) and 1 mM phenylmethanesulfonyl fluoride (Sigma-Aldrich). The suspension was incubated for 30 min at 4 °C and centrifuged at top speed in a microfuge for 15 min at 4 °C. The post nuclear lysates were collected and protein concentration was measured with the Bio-Rad Protein Assay (Bio-Rad). An equal amount of protein was mixed with 4X NuPage LDS sample buffer (Thermofisher) and 10x NuPage reducing agent (Thermofisher), warmed up at 75 °C for 5 min and loaded into NuPage Bis-Tris gels ahead of electrophoresis. The proteins were subsequently transferred to a nitrocellulose membrane (Bio-Rad and Invitrogen) via wet transfer, blocked in Tris-buffered saline containing 0.1% (v/v) Tween 20 and 5% (w/v) fetal bovine albumin for 1 h at room temperature. Immunoblots were carried out with the antibodies listed above and visualized using the ECL Western Blotting Detection Reagents (GE Healthcare). In order to strip antibodies from the blots, OneMinute Plus Western Blot Stripping Buffer (GM Biosciences) was used according to the manufacturer’s instruction. Signal was detected with Kodak’s films.

### Statistical analysis

All measurements were expressed as mean ± standard deviation. For DHT measurement of dry mass, correlation of different treatment groups was computed using a two-tailed unpaired t-test assuming unequal variances. A p-value of less than 0.05 was considered significant. The effect size was computed using Glass’s delta, which is more appropriate than Cohen’s *d* or Hedge’s *g* when the standard deviations of the compared groups are different^17^. The number of samples analyzed with Coulter counter was typically greater than 10^5^, which is large enough to reject any null hypothesis of equal distribution. Thus, we calculated only the effect size for the volume measurement. For the dry mass measurement of cell lines with 16 nM bortezomib treatment, the mass fold change of treated cells at 1 and 4 hours relative to the mass at time 0 were compared with the mass fold change of untreated cells at the corresponding time by the left-tailed unpaired t-test assuming unequal variances. A p-value of less than 0.05 was considered significant. For the dry mass measurement of patient samples, the mass change of treated and untreated samples at 1 hour and 4 hours were compared by a two-tailed unpaired t-test assuming unequal variances. A p-value of less than 0.05 was considered significant.

## Results

### Validation using HeLa cells treated with staurosporine

To validate our approach as a whole, we measured the dry mass and volume changes in HeLa cells due to the treatment with staurosporine, which is known to induce apoptosis in HeLa cells. During the process, the cell volume decreases without mass change^18^. The so-called apoptotic volume decrease (AVD) has been considered as a hallmark of apoptosis in many cell types, and is attributed to an efflux of monovalent ions and osmotic imbalance, and consequent water loss^19^. Using DHT and Coulter counter, we indeed observed that the volume of HeLa cells decreased by 10% with the mean dry mass changed only by 1.4% after 2 hours of incubation with 4 μM staurosporine (Fig. 2 and Table 1).

**Figure 2.**
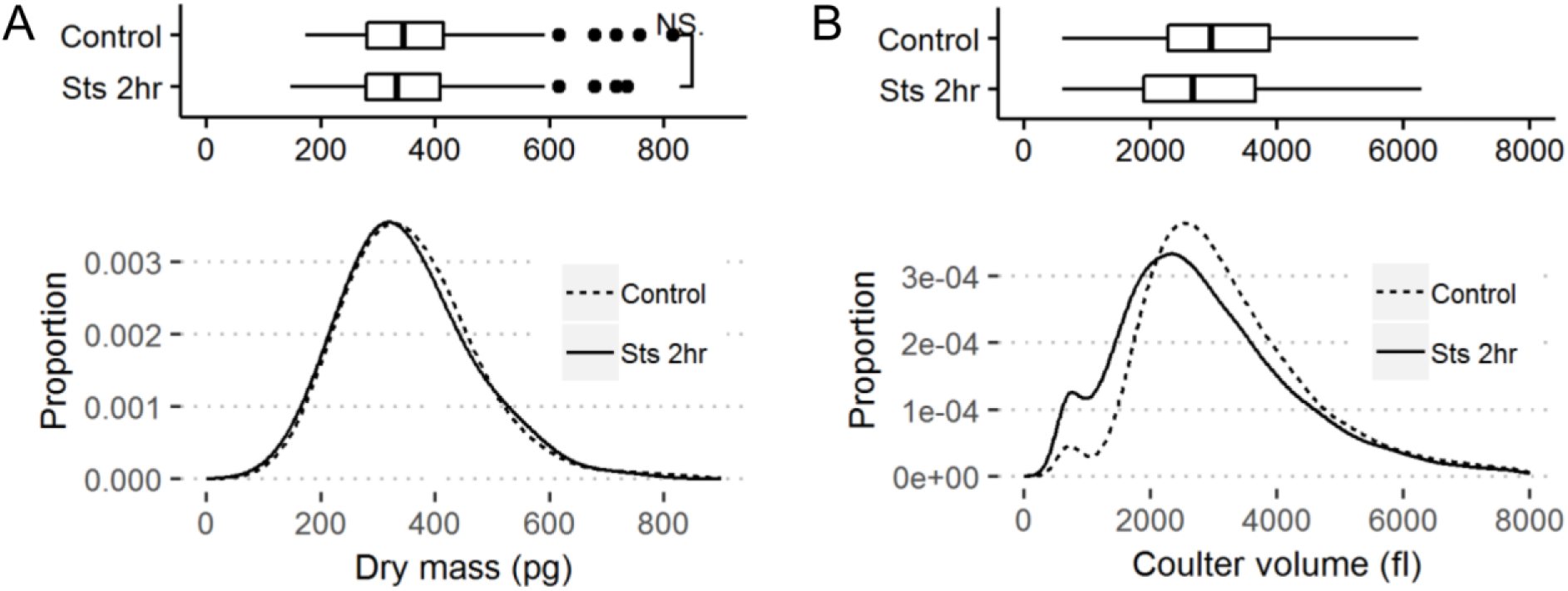
Dry mass (A) and Coulter volume (B) of the HeLa cells treated with Staurosporine (4 uM, 2 hours). A. The box and whisker plot on the top and the curve at the bottom represent the dry mass in picograms (pg) of HeLa cells treated with staurosporin at 4 μM concentration for 2 hours as compared to untreated cells. The figure represents collective data from 2 independent biological experiments. B. The box and whisker plot on the top and the curve at the bottom represent the volume in femtoliters (fl) of HeLa cells treated with staurosporin at 4 μM concentration for 2 hours as compared to untreated cells. The figure represents collective data from 2 independent biological experiments.

**Table 1.** Dry mass, volume, and Coulter volume of HeLa cells after treatment with staurosporine. The measurements are shown as mean ± standard deviation. The variables *n* and ES represent the number of samples and the effect size (Glass’s delta), respectively, and NS means not statistically significant (*p* > 0.05). The table reports aggregate data from 2 independent biological experiments.

| | Control | Staurosporine 4 μM, 2 hours |
| --- | --- | --- |
| Dry mass (pg) | 356 ± 106 ($n = 216$) | 351 ± 107 ($n = 214$)<br>ES = -0.049 (NS) |
| Coulter volume (fl) | 3276 ± 1450 ($n = 5.3 \times 10^5$) | 2942 ± 1498 ($n = 4.8 \times 10^5$)<br>ES = -0.23 |

### Dry mass and volume change of RPMI8226 cells upon treatment with bortezomib, doxorubicin, or UV irradiation

We measured the dry mass and volume of RPMI8226 cells after treatment with bortezomib (20 nM), doxorubicin (1 μM) and UV (1 min). Figure 3 and Table 2 summarize the results. When incubated with 20 nM of bortezomib, RPMI8226 cells increased dry mass by 1.1% in an hour and 12% in 4 hours (Table 2a). This dry mass increase is presumably secondary to increased proteasome load, as we will show later. Noteworthy, RPMI8226 cells showed similar dry mass increase when treated with doxorubicin and UV (Table 2b). For 1 μM of doxorubicin, RPMI8226 cells increased dry mass by 3.9% in an hour and 11% in 4 hours. Upon 1-minute UV irradiation, RPMI8226 cells increased dry mass by 2.2% in an hour and 12% in 4 hours. These similar increases in dry mass are interesting, considering the widely different mechanisms of these treatments: proteasome inhibition by bortezomib^20^, DNA intercalation by doxorubicin^21^, and DNA damage by UV irradiation^22^. However, the dry mass increase upon doxorubicin and UV treatment is not explained by the proteasome load, as we will show later.

**Figure 3.**
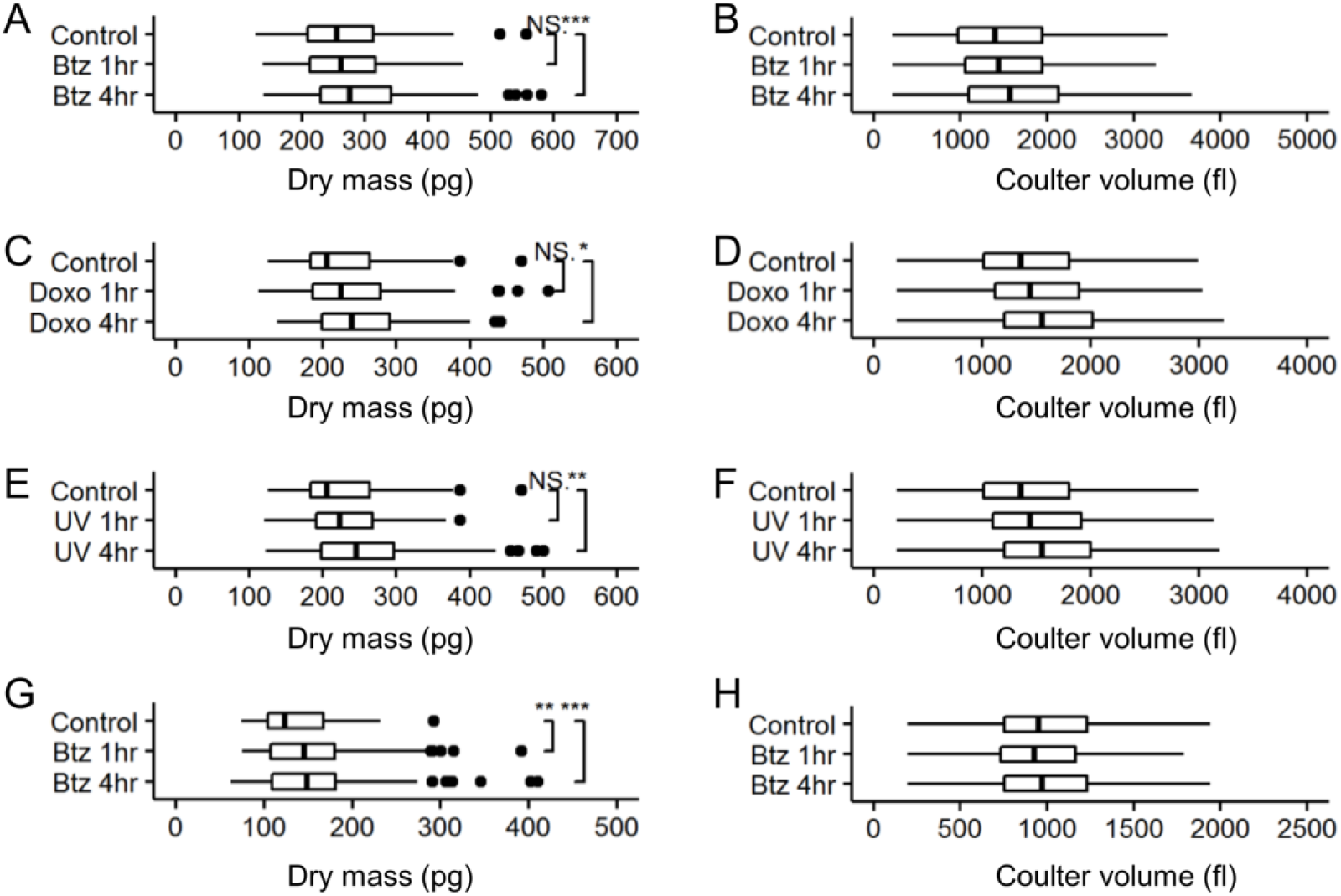
Dry mass and Coulter volume of the multiple myeloma cells treated with bortezomib (20 nM), doxorubicin (1 μM), and UV (1 min). A. The box and whisker plot represents the dry mass in picograms (pg) of RPMI8226 cells treated with Bortezomib at 20 nM concentration for 1 and 4 hours as compared to untreated cells. The figure represents collective data from 2 independent biological experiments. B. The box and whisker plot on the top and the curve at the bottom represent the Coulter volume in femtoliters (fl) of RPMI8226 cells treated with Bortezomib at 20 nM concentration for 1 and 4 hours as compared to untreated cells. The figure represents collective data from 2 independent biological experiments. C. The box and whisker plot represents the dry mass in picograms (pg) of RPMI8226 cells treated with Doxorubicin at 1 μM concentration for 1 and 4 hours as compared to untreated cells. The figure represents collective data from 2 independent biological experiments. D. The box and whisker plot on the top and the curve at the bottom represent the Coulter volume in femtoliters (fl) of RPMI8226 cells treated with Doxorubicin at 1 μM concentration for 1 and 4 hours as compared to untreated cells. The figure represents collective data from 2 independent biological experiments. E. The box and whisker plot represents the dry mass in picograms (pg) of RPMI8226 cells treated with UV for 1 minute then measured in 1 and 4 hours as compared to untreated cells. The figure represents collective data from 2 independent biological experiments. F. The box and whisker plot on the top and the curve at the bottom represent the Coulter volume in femtoliters (fl) of RPMI8226 cells treated with UV for 1 minute then measured in 1 and 4 hours as compared to untreated cells. The figure represents collective data from 2 independent biological experiments. G. The box and whisker plot represents the dry mass in picograms (pg) of MM.1S cells treated with Bortezomib at 20 nM concentration for 1 and 4 hours as compared to untreated cells. The figure represents collective data from 2 independent biological experiments. H. The box and whisker plot on the top and the curve at the bottom represent the Coulter volume in femtoliters (fl) of MM.1S cells treated with Bortezomib at 20 nM concentration for 1 and 4 hours as compared to untreated cells. The figure represents collective data from 2 independent biological experiments.

**Table 2.**
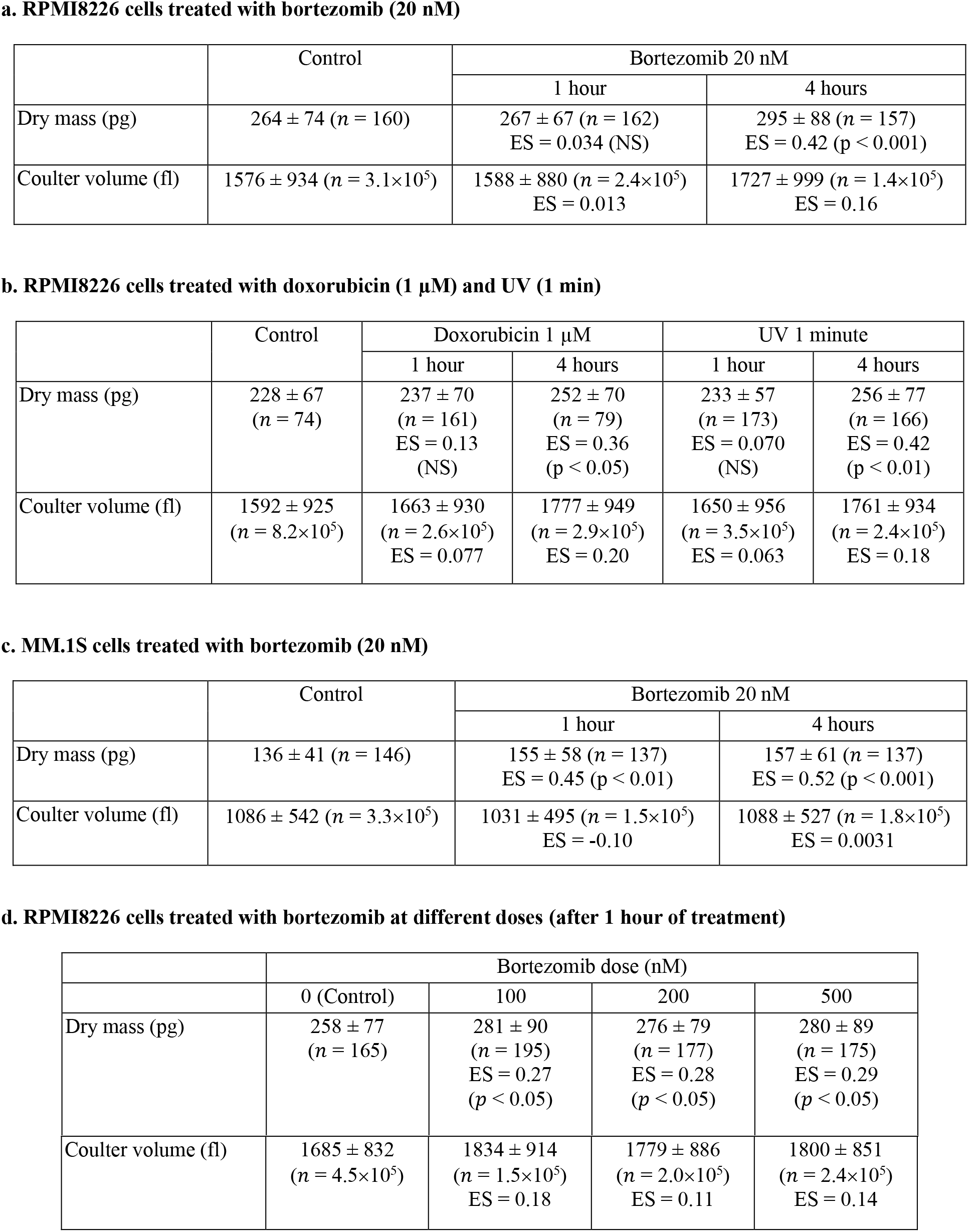
Dry mass and Coulter volume of multiple myeloma cells after treatment with bortezomib, doxorubicin, and UV. The variables *n* and ES represent the number of samples and the effect size (Glass’s delta), respectively, and NS means not statistically significant (*p* > 0.05). Each table reports aggregate data from 2 independent biological experiments.

The volume change of RPMI8226 cells followed the dry mass change; the volume increase was observed for all the treatment conditions (bortezomib, doxorubicin, and UV) we tested. As we observed with HeLa cells treated with staurosporine, many cell types are known to decrease the volume at the early stage of apoptosis without significant change in mass. The dramatic volume increase at an early stage of apoptosis, which we observed with the RPMI8226 cells, is rare but has been observed in other systems. For example, serum-deprived vascular smooth muscle cells increase their volume by about 40% in 30- to 60-min lag phase^23^. In apoptosis of HeLa cells induced by actinomycin D, volume increases of 5-30% were observed within an hour after the onset of blebbing, the time point when the volume measurement started^24^. Notably, volume kinetics in apoptotic cells can be quite complex. For example, apoptosis without cell shrinkage has been observed^25,26^. The apoptosis of Ehrlich ascites tumor cells induced by cisplatin leads to three distinctive volume change stages: an early decrease (4–12 hours), a partial recovery (12 to 32 hours), and a further reduction (past 32 hours). In contrast, the volume of MM.1S cells was reduced by 5.1% in one hour but was restored to the original value in the next three hours, i.e., four hours after the treatment (Table 2c). This volume decrease is a typical response of cells, but it contrasts with the response of RPMI8226 cells. Therefore, we use only dry mass as an indicator of the apoptosis of MM cells.

### Dry mass change of MM cells as an early indicator of bortezomib-induced apoptosis

Based on our early observation of dry mass increase in bortezomib-treated RPMI8226 cells, we checked the dry mass change of other MM.1S cells upon treatment with bortezomib. For the same treatment (20 nM bortezomib), MM.1S cells also increased the dry mass, but the response was much faster; the dry mass increased by 14% in an hour then remained at about the same level (Table 2c). This fast response may be attributed to the high drug sensitivity of MM.1S compared to RPMI8226 cells. Note that the EC_50_ of bortezomib is only 4 nM for MM.1S cells^7^, which is less than 1/10 of that for RPMI8226 cells. Indeed, when we increased the bortezomib dose to 100 nM, RPMI8226 cells increased both the dry mass (8.9%) and volume (8.8%) significantly in only one hour (Table 2d). The dry mass and volume did not change significantly when we further increased the dose to 200 and 500 nM, suggesting saturation of proteasome inhibition with 100 nM dose (Table 2d).

To evaluate whether dry mass could serve as an early biomarker of bortezomib-induced apoptosis in MM cells, we further tested KMS20 cells, AMO1 cells, and AMO1-VR (an isogenic AMO1 cell line adapted to grow in continuous bortezomib) cells to bortezomib. We quantified their sensitivity to bortezomib by the annexin V/PI apoptosis assay at 24 hours after the treatment (Fig. 4E) and chose a concentration of 16 nM, which distinguished the cell lines, for the following analysis. For the dry mass measurements, we used ceQPM with a higher imaging throughput than DHT. The dry mass measurement of MM.1S cells was repeated to confirm the consistency for the altered dose and imaging technique. Figure 4 summarizes the result. Upon the treatment with 16 nM bortezomib, the dry mass increase of MM.1S cells was not significant *(p* =0.02) after one hour; however, the increase was significant after 4 hours of treatment (Fig. 4A). Whereas, KMS20 cells, which has intermediate sensitivity to bortezomib, showed significant increase in dry mass after 4 hours (p =0.03) but with lower magnitude (Fig. 4B). In addition, AMO1 cells, which are sensitive to bortezomib, significantly increased the dry mass even in one hour after the treatment (p = 0.02) with 16 nM of bortezomib (Fig. 4C). However, AMO1-VR, which had induced resistance to bortzezomib and was the most insensitive cell line in our apoptosis assay, showed the least dry mass increase after 4 hours (Fig. 4C). Moreover, we found a strikingly good correlation between the mass fold change of treated cells at 4 hours relative to time 0 and the percentage of dead cells at 24 hours, suggesting a quantitative relation between the rapid mass accumulation and apoptosis caused by bortezomib.

**Figure 4.**
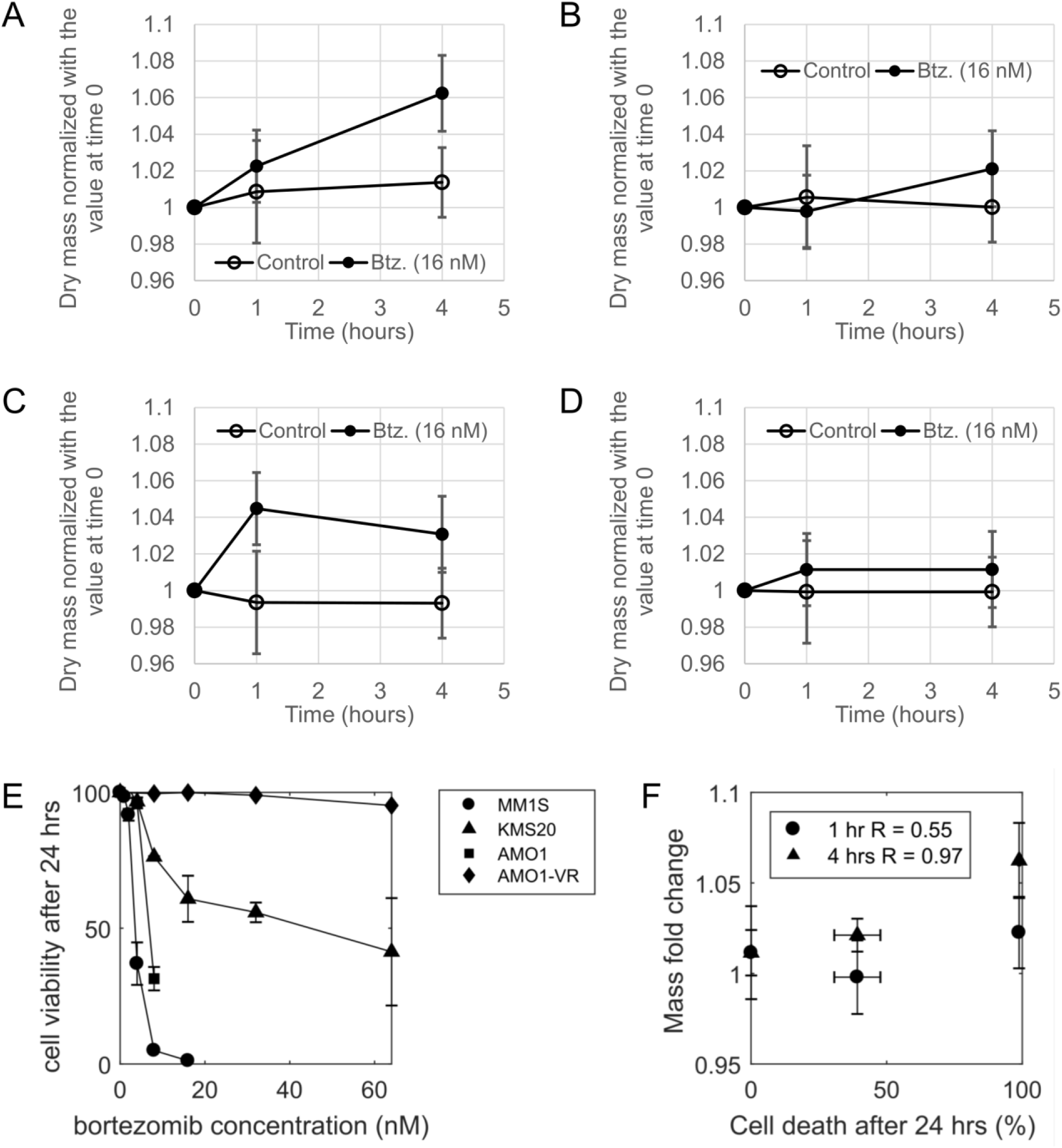
Dry mass change after the treatment with 16 nM bortezomib in (A) MM.1S, (B) KMS20, (C) AMO1, and (D) AMO1-VR cell lines. <u>Error bars indicate the standard deviation of three independent experiments</u>. (E) Cell viability at 24 hours after bortezomib treatment at indicated concentrations. Error bars are the standard deviation of replicated experiments. (F) Correlation between mass fold changes of treated cells at 1 or 4 hours with the percentages of cell death at 24 hours after 16 nM bortezomib treatment. Error bars are the standard deviation of the measurements derived from three independent experiments. R is the Pearson’s correlation.

### Dry mass increase 4 hours post bortezomimb treatment positively correlates with bortezomib sensitivity in primary MM cells from patients

Finally, we performed the dry mass measurement on primary MM cells derived from four patients, three newly-diagnosed, treatment naive and one relapsed-refractory patients previously exposed to bortezomib. Similarly, to the experiments with MM cell lines, we treated cells for 1 or 4 hours with 16 nM bortezomib prior to measuring the dry mass, as previously described. In parallel, we treated primary MM cells for 24 hours with 16 nM bortezomib and assess apoptosis via annexin V/propidium iodine staining and flow cytometry analysis. Figure 5 summarizes the result. Interestingly, across the treatment naïve, MM samples, NDMM2 showed no significant increase in dry mass at 1-hour post treatment and a reduction in dry mass at 4 hours post treatment with bortezomib. Across all samples analyzed, this sample showed the least sensitivity to bortezomib, suggesting innate resistance. In fact, the dry mass change measured after 4 hours of bortezomib treatment (16 nM) strongly correlated with the ratio of apoptotic cells measured after 24 hours. All these data using additional cell lines and primary cells confirm our observation that the dry mass increase could serve as an early marker for the bortezomib-induced apoptosis of MM cells.

**Figure 5.**
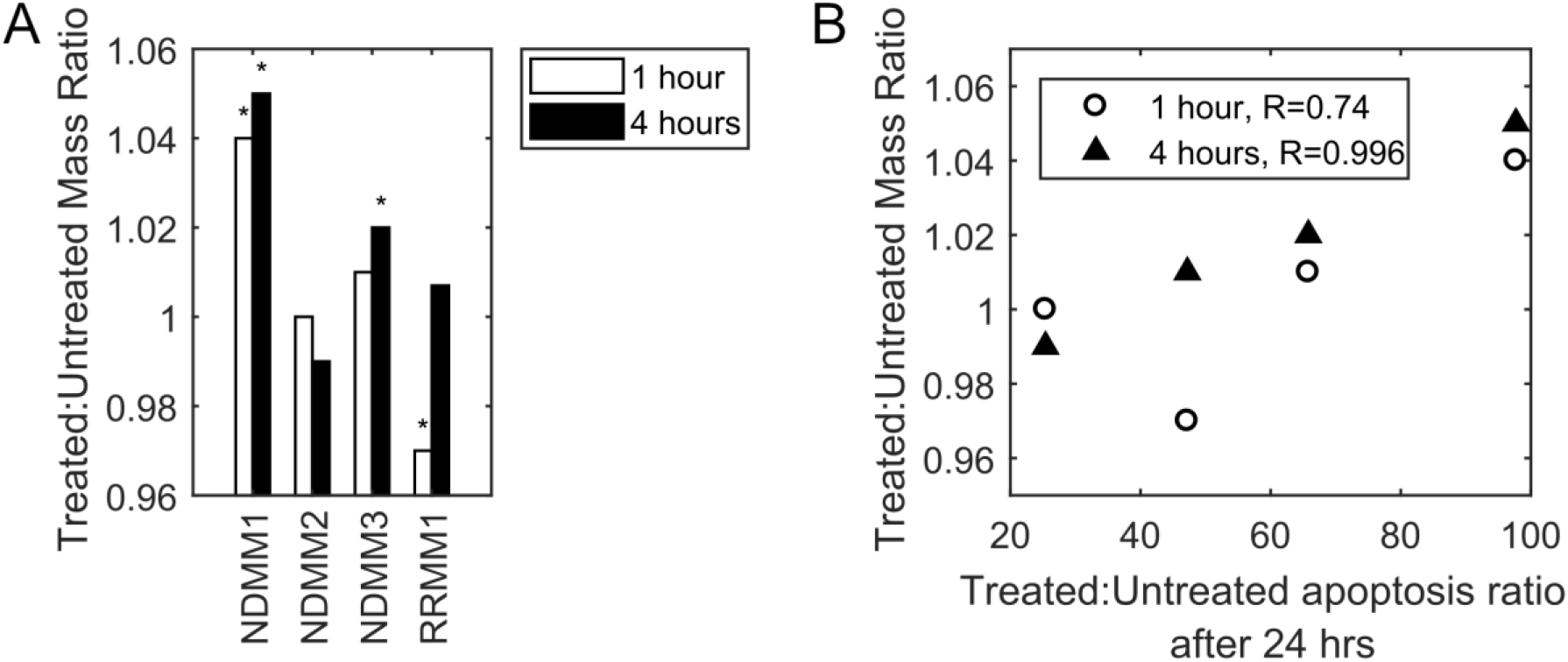
Dry mass change in four primary MM samples from patients: (A) the dry mass ratio of bortezomib-treated versus untreated cells, and (B) the values plotted with the apoptosis percentage ratio of treated versus untreated, as measured after 24 hours of treatment. NDMM: newly-diagnosed MM; and RRMM: relapsed/refractory MM. (*) in (A) indicates significant difference (p<0.05) between the treated and untreated samples. R in (B) is the Pearson’s correlation. For each test case, we imaged about 70 fields-of-view (FOV) within 40 minutes. Each FOV typically contains 1-2 cells.

### Dry mass changes are cell cycle phase independent

Cellular dry mass distribution can be affected by cell cycle distribution since G2-M phase cells are larger than G1 phase cells; cell cycle arrest at mitosis may shift the cell size distribution to larger average size. To check whether the observed dry mass and volume changes are due to such cell cycle arrest, we obtained the cell cycle distributions of RPMI8226 cells for different treatments used in this study (Fig. 6). With bortezomib, there was no significant change in the cell cycle distribution. With doxorubicin and UV exposure, the G1 population slightly increased, which would decrease the dry mass of cell population instead of increasing it as we observed. Then, the observed changes in treated cells can be attributed to the response of individual cells irrespective of their positions on the cell cycle. This cell cycle-independent response is important, since it means that we can assess the effect of a certain treatment by monitoring the responses of a small number of cells in an asynchronous population.

**Figure 6.**
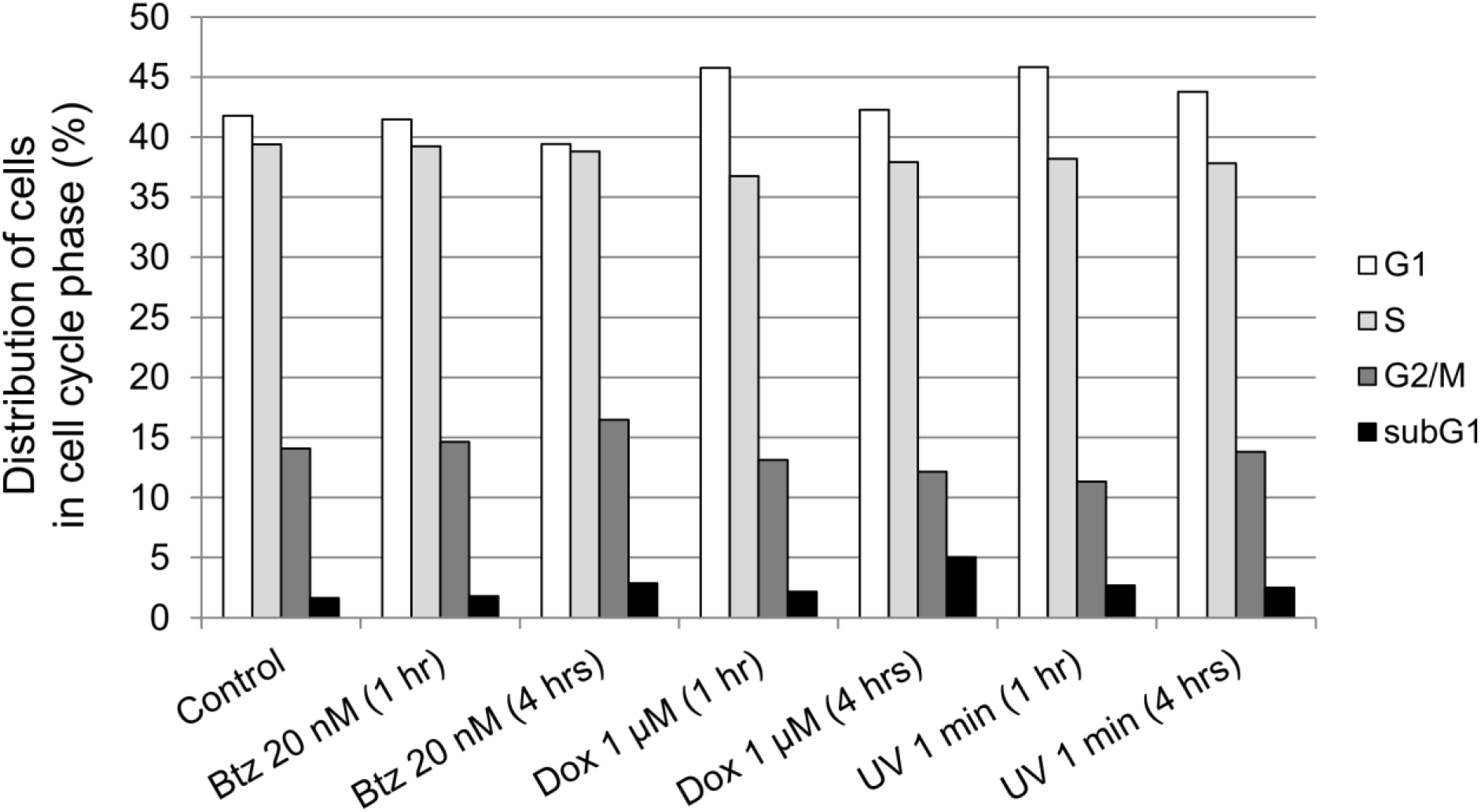
Cell cycle analysis. Cell cycle analysis of RPMI cells treated as indicated shows no significant changes compared to control cells in the proportion of cells arrested in S phase, G2/M or in apoptosis (subG1) after treatment as indicated. One of three independent experiments is shown.

### Dry mass changes in MM cells upon treatment precede validated markers of apoptosis

Importantly, currently available assays are not sensitive enough to detect early commitment to apoptosis induced by these agents. For example, Fig. 7A shows a flow cytometry analysis with annexin V/PI staining to detect apoptosis in RPMI cells after the indicated treatment. Annexin V binds avidly to the phospholipid phosphatidylserine (PS) that is translocated from the inner to the outer leaflet of the plasma membrane in the early stages of apoptosis. Therefore, annexin V positivity in non-permeabilized cells, reflects PS exposure extracellularly and is a validated marker of early apoptosis. Instead, propidium iodine binds to DNA and positivity to this markers signal that the intracellular DNA is freely accessible, suggesting loss of membrane integrity, a late stage of apoptosis. The percentage of viable cells was not significantly changed at the end of any of the treatments we performed (black bars). Noteworthy, when the drug was washed out and cells were re-plated for 20-23 hours (grey bars), apoptosis was detectable on flow cytometry and via PARP cleavage in RPMI cells treated for 4 hours with bortezomib 20nM or for one hour with 100-500nM bortezomib, suggesting that cells had committed to apoptosis (Fig. 7A and 7B). In contrast, the UV-treated cells did not show any loss of cell viability or cleavage of PARP when allowed to grow for 24 hours post irradiation, suggesting that exposure length and intensity was not sufficient to commit cells to apoptosis.

**Figure 7.**
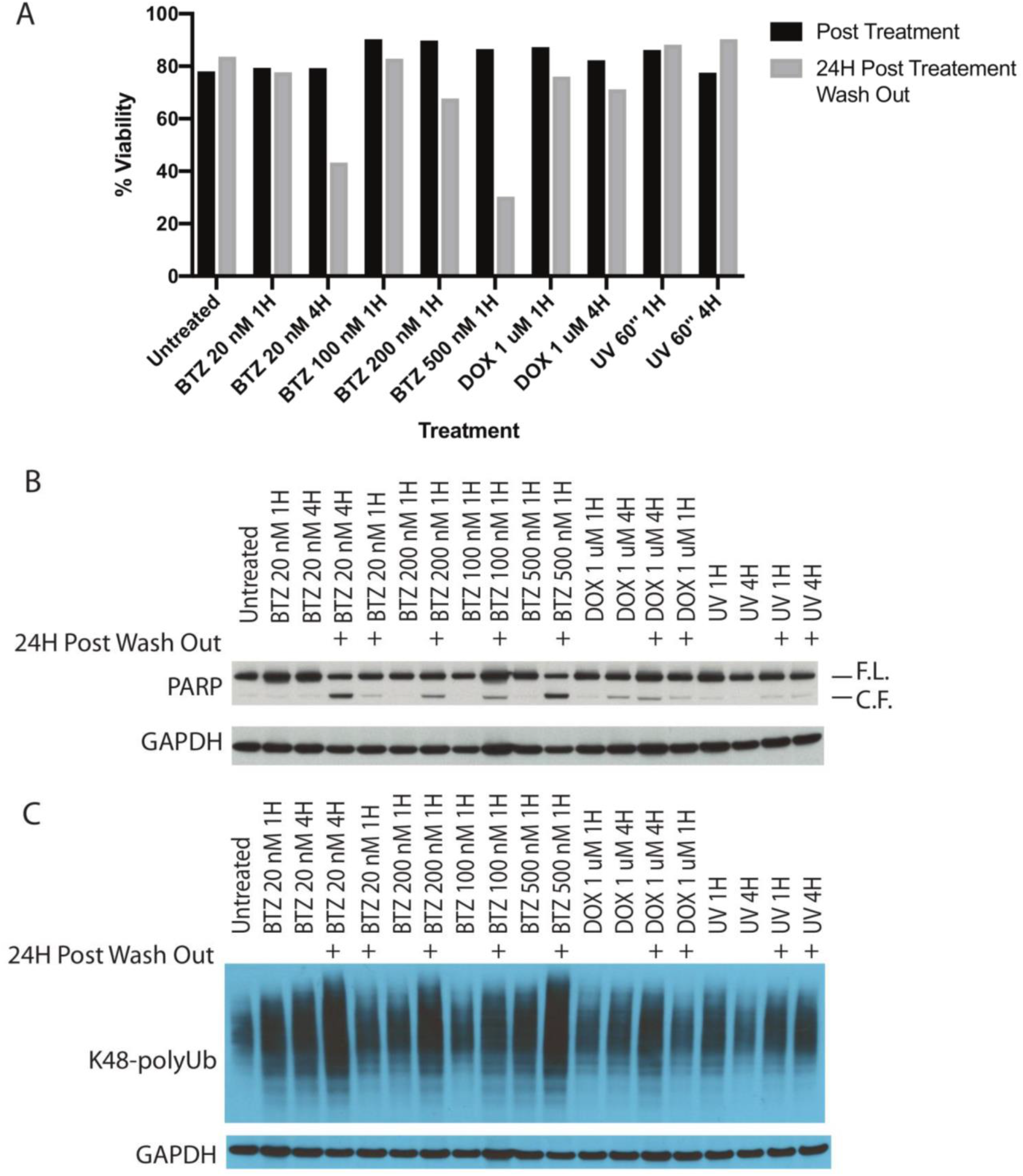
Apoptosis and K48-polyubiquitin accumulation in RPMI8226 cells. (A) Flow cytometry analysis with annexin V/PI staining of RPMI cells harvested right after indicated treatment (black bars) or after a total of 24 hours wash out period from treatment (grey bars). The y axis shows absolute percentage of viable (double negative, annexin V/PI) cells. One of three independentent experiments is shown. (B) Western blotting analysis of PARP cleavage, a signature of apoptosis, in RPMI8226 cells harvested right after indicated treatment or after a total of 24 hours wash out period from treatment. Consistent with flow cytometry data, 4 hour treatment with 20nM bortezomib or 1 hour treatment with 200 and 500 nM bortezomib results in commitment to apoptosis and appearance of PARP cleaved fragment (C.F.) after drug wash out. One of three independentent experiments is shown. (C) Western blotting analysis of K48 polyubiquitin protein accumulation. The full-length images of the blots shown in (B) and (C) are included as Supplementary Figures 1 and 2. One of three independentent experiments is shown.

These data are consistent with western blotting analysis for Poly (ADP-ribose) polymerase (PARP) cleavage, a marker of apoptosis, after indicated treatments. When cells were harvested at the end of treatment (lanes without ‘+’ mark on 24H Post Wash Out), PARP cleavage was not significantly increased across all treatments examined, except for a modest increase in cells treated with 1 μM of doxorubicin for 4 hours.

To assess whether accumulation of polyubiquitinated proteins could explain the dry mass increase in bortezomib-treated cells, we assessed K-48-linked polyubiquitinated (polyUb) proteins after the indicated treatments with or without drug washout and re-plating to complete 24 hours. Consistent with our prior data, treatment with bortezomib leads to increased K-48-linked polyUb proteins (Fig. 7C). However, the dry mass increase in doxorubicin- or UV-treated cells is not fully explained by the proteasome workload, as the increase of K48-polyubiquitinated proteins in these cells is much less significant than in bortezomib-treated cells (Fig. 7C).

## Discussion

In this study we provide proof of concept of feasibility and potential utility of measuring dynamic changes in cellular dry mass to predict response to proteotoxic and genotoxic treatments in MM cell lines and primary cells from patients. In particular, we focused on evaluating cellular dry mass upon treatment with the proteasome inhibitor bortezomib across cell lines and primary cells with distinct sensitivity. Overall, our data in cell lines suggest that an increase in cellular dry mass as early as 1-4 hours post treatment with bortezomib may be a useful early biomarker of sensitivity. Further, across a small sample of primary MM cell lines, we were able to observe a tight correlation between extent of increase in cellular dry mass post 4 hours of bortezomib treatment and sensitivity to bortezomib as measured via apoptosis. Importantly, the increase in dry mass precedes the appearance of typical signatures of apoptosis, e.g., bleb formation, caspase activity increase, PARP cleavage, membrane translocation^27^.

Of note, this approach proved feasible in primary samples of variable cellularity obtained from MM patients via bone marrow aspirate through standard density gradient and magnetic bead isolation.

Our data could not completely confirm our initial hypothesis that accumulation of polyUb proteins is at the base of dry mass increase and predictive of bortezomib sensitivity. In fact, while we observed polyUb accumulation in samples treated with bortezomib, treatments that did not trigger polyUb accumulation, such as doxorubicin and UV, resulted in a similar increase in dry mass, preceding apoptosis. This data suggest that mechanisms other than polyUb accumulation are likely underlying dry mass changes upon drug treatment. We are planning to investigate the underlying mechanisms of dry mass increase in our future work.

Of note, while we were unable to correlate dry mass changes with specific subclones of distinct genomic background, we observed consistent changes across the entire distribution of single cells analyzed, presumably encompassing genomically heterogeneous cells. Furthermore, our observations were reproducible on 4 MM cell lines characterized by different genetic mutations: RPMI is characterized by t(16;22) and t(8;22), G12A heterozygous mutation in KRAS, and a E285K homozygous mutation in TP53; MM1S, harbors a t(14:16) and a t(8;14), a heterozygous G12A KRAS mutation and is TP53 wild type (WT); KMS20 carries a G12S homozygous mutation in KRAS and a Y126X homozygous mutation in TP53; AMO1 harbors a t(12;14), heterozygous A146T KRAS mutation and is WT for TP53.

Our study has several limitations: first, only a limited panel of MM cell lines and a small pool of primary samples were analyzed. While this can be sufficient to provide proof of concept of feasibility and potential utility, a larger number of samples is essential to confirm our initial observation. Ideally, prospective collection of samples and correlation with clinical response to treatment and underlying genomic background could be extremely valuable to examine the potential use of this tool in the clinics. Second, we did not evaluate the reliability of this method when assessing for response to combinatorial treatments. As previously mentioned, MM induction therapy makes use of combination of drugs, often three, sometimes four distinct compounds that are administered concomitantly to patients to debulk disease. While there is certainly value in determining intrinsic sensitivity/resistance to single chemotherapy agents, the implementation of this tool to predict response to combinatorial treatment would be helpful. However, currently available combinatorial treatments boost overall response rates that are approaching 100% and identifying truly primary refractory patients would require analysis of a substantial number of patients. Third, we began this study by using digital holographic tomography (DHT), a highly accurate, but difficult to apply method of measuring dry mass. Subsequently, we confirmed our initial data by using scalable and high throughput computationally-enhanced quantitative phase microscopy (ceQPM). This methodology could potentially be applicable and reproducible across centers with adequate training, however, intrinsic expertise is required in the analysis of data, suggesting that translation of this potential tool to the clinics would require significant effort. Finally, the mechanisms driving dry mass increase upon proteotoxic and genetoxic treatments are only partially understood and it would be of critical importance to understand these determinants to derive appropriate conclusions.

Overall, we believe that these data provide proof of concept that measurement of dry mass on a single cell level could be of relevance as a sensitive predictive biomarker of response to drug treatment in MM with potential, future applications to further improve the personalized care of MM patients and maximize the chances of selecting a highly effective induction treatment on a patient-per-patient basis.

## Acknowledgments

The authors acknowledge the late professor Michael Feld for initiating this research. This work was funded by the National Institutes of Health (P41EB015871-33, R01GM026875, R21GM135848), the National Science Foundation (DBI-0754339), and Hamamatsu Photonics, Japan. This work was in part funded by the National Institute of Health/National Cancer Institute (5K08CA245100, GB). XL and SO were supported by the National Institute of Health (R56AG073341). Some early findings reported in this article were published in the Ph.D. dissertation (MIT, 2011) by Yongjin Sung. The authors thank Professor Marc Kirschner of Harvard Medical School for making available the ceQPM microscope and analytical tools.

## Author Contributions

W.C., K.C.A. G.B. and Y.S. developed the concept; X.L., W.C., G.B. and Y.S.. designed the experiments; X.L., M.M., S.O., T.C., B.E., G.B. and Y.S. performed the experiments; X.L., M.M., S.O., W.C., T.C., B.E., Z.Y. G.B. and Y.S. analyzed the data; S.M.R., O.N., C.C.M., A.S.S. and G.B. contributed primary samples; Z.Y. provided technical advice; Z.Y., G.B. and Y.S. secured funding for research; X.L, G.B. and Y.S. wrote the manuscript; all authors read and approved the final version of the manuscript.

