## Supplementary Figures for "Characterizing dry mass and volume changes in human multiple myeloma cells upon treatment with proteotoxic and genotoxic drugs" (from another version of this preprint: version 1, 2022-03-19)

**Supplementary Figure 1**

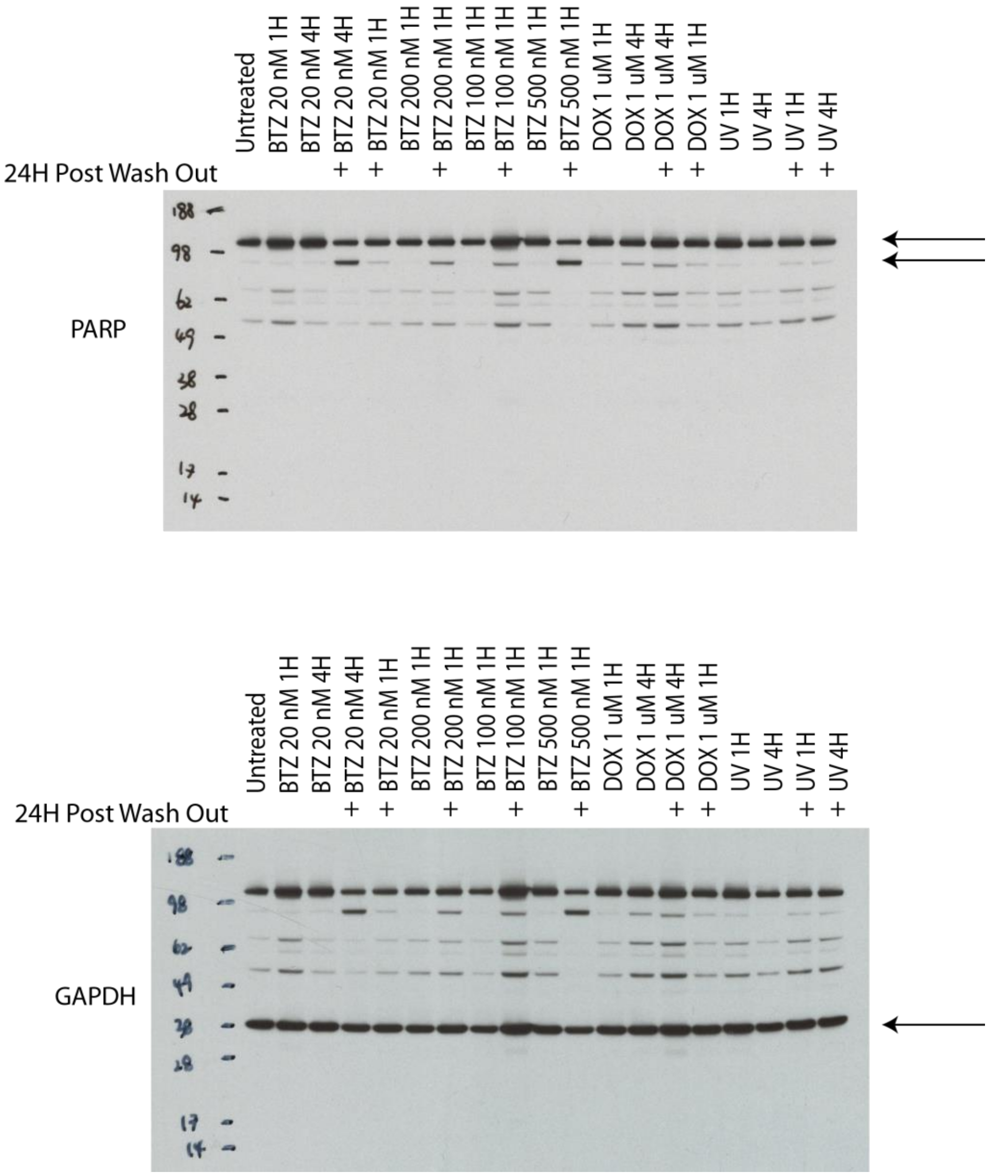


**Supplementary Figure 1.** Whole western blot membrane of PARP and GAPDH cropped images. Arrows indicate band(s) of interest. Molecular weight reference is annotated on the left side.

**Supplementary Figure 2.**

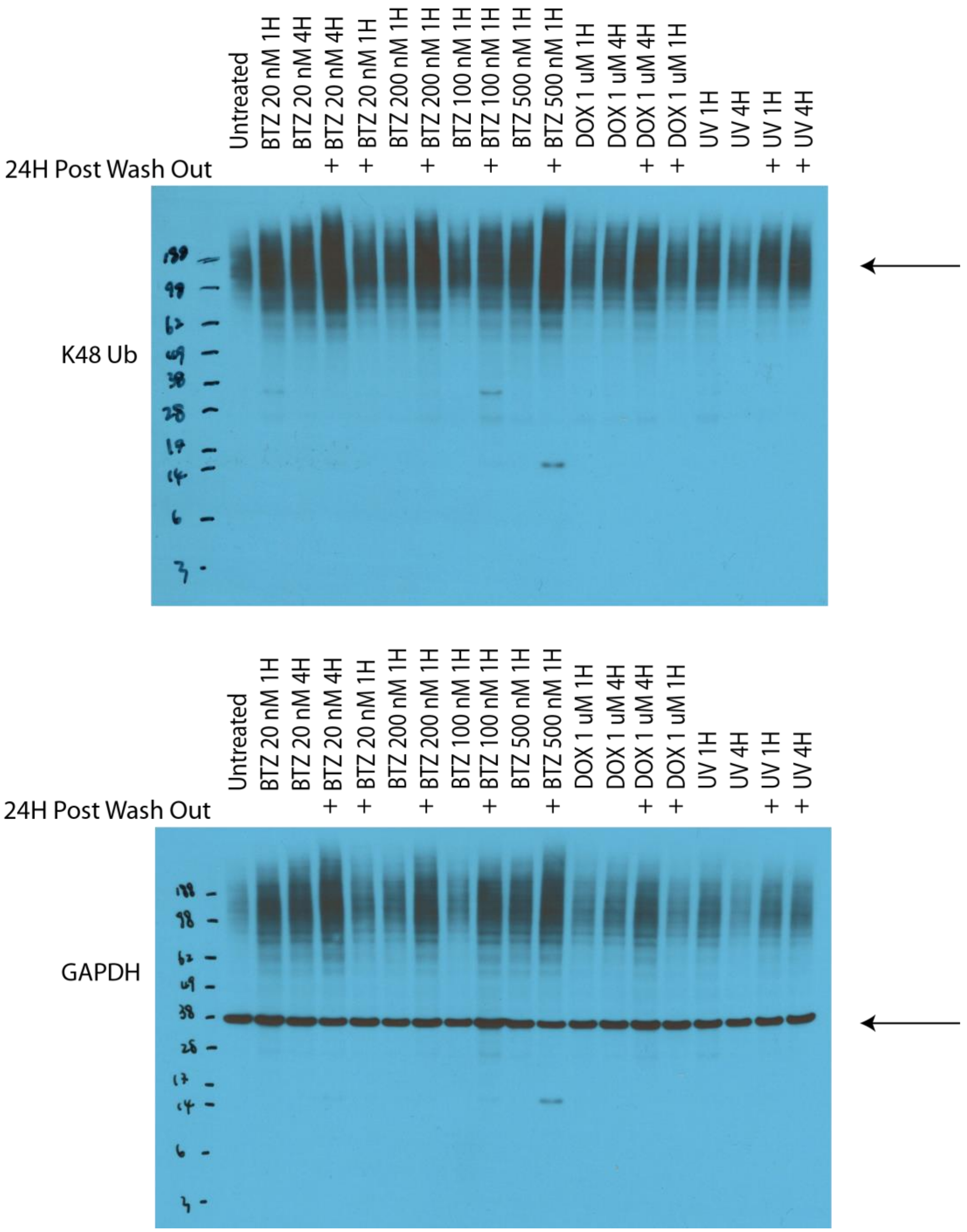


**Supplementary Figure 2.** Whole western blot membrane for K48-ubiquitin and GAPDH cropped images. Arrows indicate band(s) of interest. Molecular weight reference is annotated on the left side.
